# Unifying Temporal Phenomena in Human Visual Cortex

**DOI:** 10.1101/108639

**Authors:** Jingyang Zhou, Noah C. Benson, Kendrick Kay, Jonathan Winawer

## Abstract

Neuronal responses in visual cortex show a diversity of complex temporal properties. These properties include sub-additive temporal summation, response reduction with repeated or sustained stimuli (adaptation), and slower dynamics at low stimulus contrast. Here, we hypothesize that these seemingly disparate effects can be explained by a single, shared computational mechanism. We propose a model consisting of a linear stage, followed by history-dependent gain control. The model accounts for these various temporal phenomena, tested against an unusually diverse set of measurements - intracranial electrodes in patients, fMRI, and macaque single unit spiking. The model further enables us to uncover a systematic and rich variety of temporal encoding strategies across visual cortex: First, temporal receptive field shape differs both across and within visual field maps. Second, later visual areas show more rapid and pronounced adaptation. Our study provides a new framework to understand the transformation between visual input and dynamical cortical responses.

**Author Summary:** The nervous system extracts meaning from the distribution of light over space and time. Spatial vision has been a highly successful research area, and the spatial receptive field has served as a fundamental and unifying concept that spans perception, computation, and physiology. While there has also been a large interest in temporal vision, the temporal domain has lagged the spatial domain in terms of quantitative models of how signals are transformed across the visual hierarchy. Here we present a model of temporal dynamics of neuronal responses in human cerebral cortex. We show that the model can accurately predict responses at the millisecond scale using intracortical electrodes in patient volunteers, and that the same model generalizes to multiple types of other measurements, including functional MRI and action potentials from monkey cortex. Further, we show that a single model can account for a variety of temporal phenomena, including short-term adaptation and slower dynamics at low stimulus contrast. By developing a computational model and showing that it successfully generalizes across measurement types, cortical areas, and stimuli, we provide new insights into how time-varying images are encoded and transformed into dynamic cortical responses.

## Introduction

Sensory systems are confronted with the challenge of extracting behaviorally relevant information from the large quantity of inputs spread over space and time. In the visual system, prioritizing some inputs over others begins at the earliest stages of processing. For example, center-surround receptive fields enhance sensitivity to contrast, while attenuating sensitivity to diffuse illumination (1). And nonlinearities such as spike thresholding reduce the redundancies found in natural images (2). As a result of such processes, the representation of images is substantially transformed along the visual pathways to accentuate the importance of particular properties of the inputs for later visual processing (3).

In the visual temporal domain, a number of non-linear neuronal phenomena have also been observed, suggesting coding strategies that prioritize some temporal patterns over others. First, the neuronal response to a sustained stimulus gradually declines following an initial transient, (e.g., 4, 5) (Figure 1A). Second, responses to longer stimuli are less than the linear prediction from briefer stimuli (5, 6) (Figure 1B). Third, when two stimuli are presented close in time, the response to the second stimulus is reduced compared to the first (4, 6, 7) (Figure 1C). Fourth, the dynamics of neuronal responses depends on stimulus contrast - compared to the response to higher contrast stimuli, the response to lower contrast stimuli exhibits a phase delay, together with an amplitude reduction (6, 8, 9) (Figure 1D). These phenomena are consistent with the idea that new inputs and more reliable inputs (higher contrast) are given more weight for later processing.

**Figure 1.**
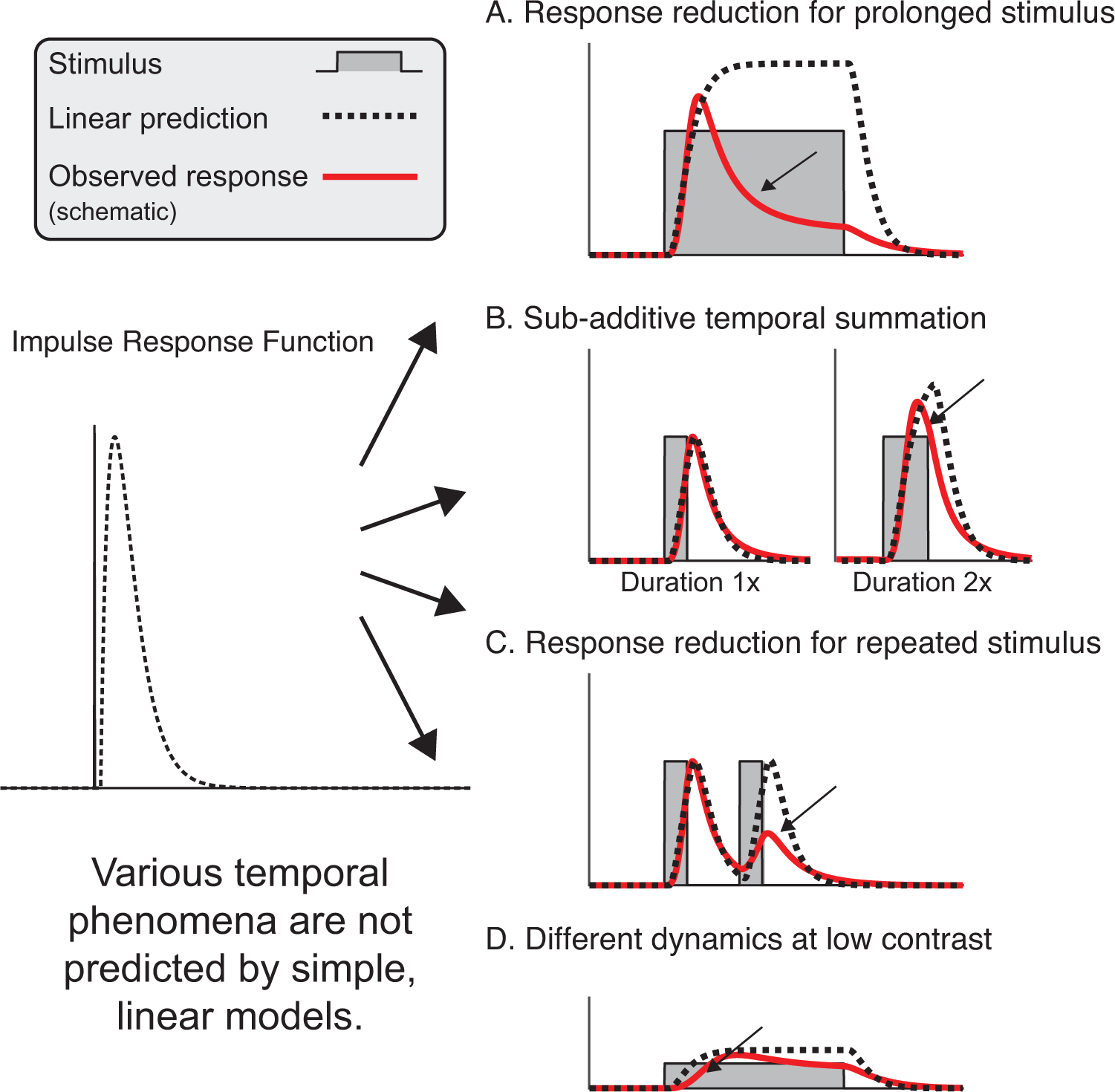
Linear model fails to predict a variety of temporal phenomena. Each panel, A-D, demonstrates temporal phenomena that have been previously been observed. For each phenomenon, we show a schematic with a stimulus time course (gray), a linear prediction (black dashed line), and a hypothetical (non-linear) observed time course (red line). The linear prediction is the result of convolving an impulse response function (left) with the stimulus time course. The hypothetical non-linear line is similar to those from prior observations. A. The neuronal response to a sustained stimulus reduces after an initial transient, differing from the linear prediction. B. The neuronal response shows subadditive temporal summation: doubling the stimulus duration results in a lower response than predicted from the linear model. C. The response to a brief stimulus is lower when there are there are two stimuli with a short gap between them. D. The response to a stimulus with lower contrast is delayed and has less of an onset transient compared to the response with high contrast (e.g., Panel A), violating linearity.

While the existence of these various temporal phenomena is well established, what is missing is a unified understanding of the neural computations that give rise to them. A central goal in sensory neuroscience is to achieve such an understanding through the development of general models that predict a variety of empirical observations (10–12). Here, we demonstrate a general model of temporal processing that takes an arbitrary stimulus time course as input and predicts the neuronal response as output. The goals in developing and testing such a model are to ask whether (1) we can provide a unified account for the seemingly disparate temporal phenomena previously observed, (2) we can leverage the model to describe systematic changes in temporal response properties across the visual hierarchy, and (3) the same model can predict responses measured with different instruments at different spatial and temporal scales.

## Results

### Dynamic normalization model: its form and its predicted dynamics

The model we present is comprised of canonical neuronal computations, with a linear first stage and a non-linear second stage (Figure 2A). The model convolves the stimulus time course with an impulse response function and then performs divisive normalization. Critically, the normalization depends on the recent history of the linear output (implemented by a low-pass filter), causing the normalization to be sluggish compared to the linear filtering. As a result, the initial response is large (relatively unnormalized) and then declines gradually as the influence of normalization increases. The idea of a divisive normalization signal that depends on response history was theorized as part of a feedback circuit model (13, 14), which was proposed to show how the steady state normalization equation might result from network activity. More commonly, normalization is implemented in a way that depends only on the instantaneous response (6, 9, 15, 16). Because in our model, the normalization depends on response history, and because this dependency is essential for much of the important model behavior, we refer to it as a dynamic normalization (DN) model.

**Figure 2.**
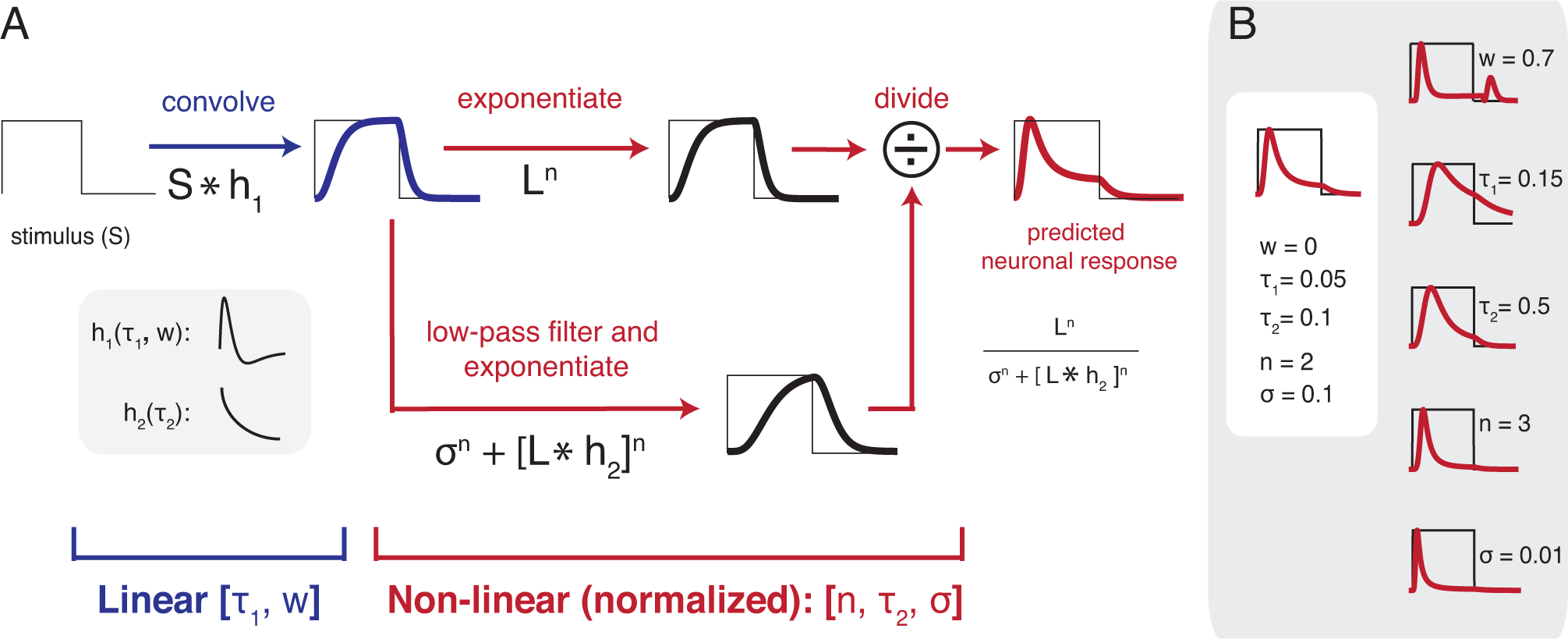
The Dynamic Normalization (DN) model. *(A)* The input to the model is the stimulus time course, S, which is 0 when the stimulus is absent and 1 when it is present. The first model stage is linear, computed by convolving S with an impulse response function, *h*_1_ (parameterized by τ_1_ and *w).* The second stage is nonlinear. The nonlinearity includes exponentiation and divisive normalization. The numerator is the linear response time course raised point-wise to a power *n*, assumed to be greater than 1. This predicted response is then divisively normalized, with the normalization being the sum of a semi-saturation constant (*σ*), and a low-pass filtered linear response (parameterized by time constant τ_2_), each raised to the same power *n*. The predicted neuronal response (red curve, right) to the example input (gray curve, left) includes a transient followed by a lower-level, more sustained response. (B) The effects of varying each of the 5 parameters are shown. For example, when *w* is increased (top) the IRF is biphasic resulting in a large response following stimulus offset. In each simulation, the parameters are *w*=0, τ_1_=0.05, τ_2_=0.1, n=2, σ=1, except for the labeled parameter.

The DN model is parameterized by five variables: τ_1_, τ_2_, *w, n* and *σ.* In the linear computation, the impulse response function (IRF) is the weighted difference between two gamma functions, similar to the IRF used by Watson (17), with the weight (*w*) between 0 and 1 on the second (negative) gamma function. A large *w* results in a large transient response at both stimulus onset and offset (Figure 2B). When fitting the DN model to time-resolved data, unless specified otherwise, we fix *w* to be 0 to reduce the number of free parameters and because the offset transient response is small in most data. The exception is in the periphery, where the offset response is larger. Hence when modeling individual cortical locations, we allow *w* to vary. The second variable that parameterizes the linear computation is a time constant τ_1_, the time to peak in the IRF. T_1_ controls the width of the impulse response, therefore the length of temporal summation in the DN model. The remaining three variables parameterize the history-dependent divisive normalization. The numerator of the normalization computation is the linear response raised point-wise to some power *n.* When *n* is 2, this is called an energy calculation (18, 19). The denominator has two terms, a semi-saturation constant, σ, and a low-passed linear response, parameterized by time constant τ_2_. Each term is raised to the same power *n*. In some models of normalization, the term in the numerator is thought of as the linear response of a cell, and the term in the denominator as the pooled response of the neighboring cells. Here, we assume the same time course for the linear terms in both the numerator and the denominator. This would be expected from modeling the sum of a population of neurons with a shared normalization pool, as shown previously (20). A generalization of the model, not implemented here, would allow for distinct time courses in the numerator (e.g., the response of a single cell or neuronal sub-population) and denominator (the normalization pool, comprised of a larger population).

Following stimulus onset, the DN model output increases rapidly due to convolution and exponentiation, and then reduces due to normalization, remaining at a lower, sustained level until stimulus offset. Although summation (convolution) and adaptation (normalization) both occur continuously throughout the predicted time course, different parts of the time course emphasize different neuronal phenomena: The initial response increase primarily reflects temporal summation (combining current inputs with past inputs), whereas the reduction following the initial transient demonstrates adaptation, since the response level declines when the stimulus is unchanging.

In the remaining parts of the Results, we used data from different measurement techniques to examine the 4 temporal phenomena shown in Figure 1: reduced responses for prolonged stimuli (ECoG), subadditive temporal summation (fMRI), reduced responses for repeated stimuli (fMRI), and delayed response at low contrast (single unit spike rate). Because the stimuli differed across experiments, different datasets exhibit different phenomena (for example, the BOLD experiment varied the stimulus duration but not the contrast, and the single unit experiments varied the contrast but not duration).

### Phenomenon 1: Response reduction for prolonged stimuli

In this section, we show that the DN model captures the general shape of neuronal responses measured using different methods, in particular showing that for static images of a few hundred ms, the model accurately predicts the initial transient followed by the reduced sustained response (Figure 1A). Further we show how the details of this transient/sustained pattern differ across cortical locations, and how these different patterns correspond to differences in the DN model parameters.

### Differences along the visual hierarchy

First, we extracted the envelope of the high frequency (70-210 Hz, ‘broadband’) time courses from a large set of human ECoG electrodes spanning multiple visual field maps. The spectral patterns in these electrode responses (but not the time courses) were analyzed for a prior publication (21). We binned the electrodes into four ROIs (V1, V2, V3, and anterior maps) based on their cortical locations and estimated receptive field centers from a separate retinotopy experiment. The “anterior maps” ROI includes electrodes from ventral (hV4, VO-1/2), lateral (LO-1/2), and dorsal (V3A/B, IPS) visual field maps. They were binned into one ROI to match the number of electrodes in the V1-V3 ROIs (n=12, 15, 11, 12; V1, V2, V3, anterior). In each trial during the experiment, a static texture (22°-diameter) was presented for 500 ms followed by a 500-ms blank. The textures were noise patterns with *1/f*^*n*^ amplitude spectra, and *n* = 0, 1, or 2 (white, pink, or brown noise). The experiment also included large field grating stimuli, but these were not included for analysis because they elicit unusual time courses (large, narrowband gamma oscillations). We averaged the broadband time series across stimulus class, trials, and electrodes within each ROI before fitting the average time series with the DN model. (See Figure S1 for individual electrode locations and responses).

The DN model provided an excellent fit to the broadband time course from all 4 ROIs, with the variance explained by the model between 90% and 99% (Figure 3). The responses in each of the 4 ROIs exhibited the characteristic pattern whereby the amplitude substantially declined following an initial large response (e.g., as depicted in the schematic in Figure 1A). The largest amplitude responses were in the earliest areas: from 7-fold over baseline in V1 to ~1.5-fold in the anterior maps. In addition to amplitude differences, there were also quantitative differences in the shape of the time courses from different ROIs. These differences were reflected in both the model predictions and in summary metrics derived from the model fit (Figure 3, right panel).

**Figure 3.**
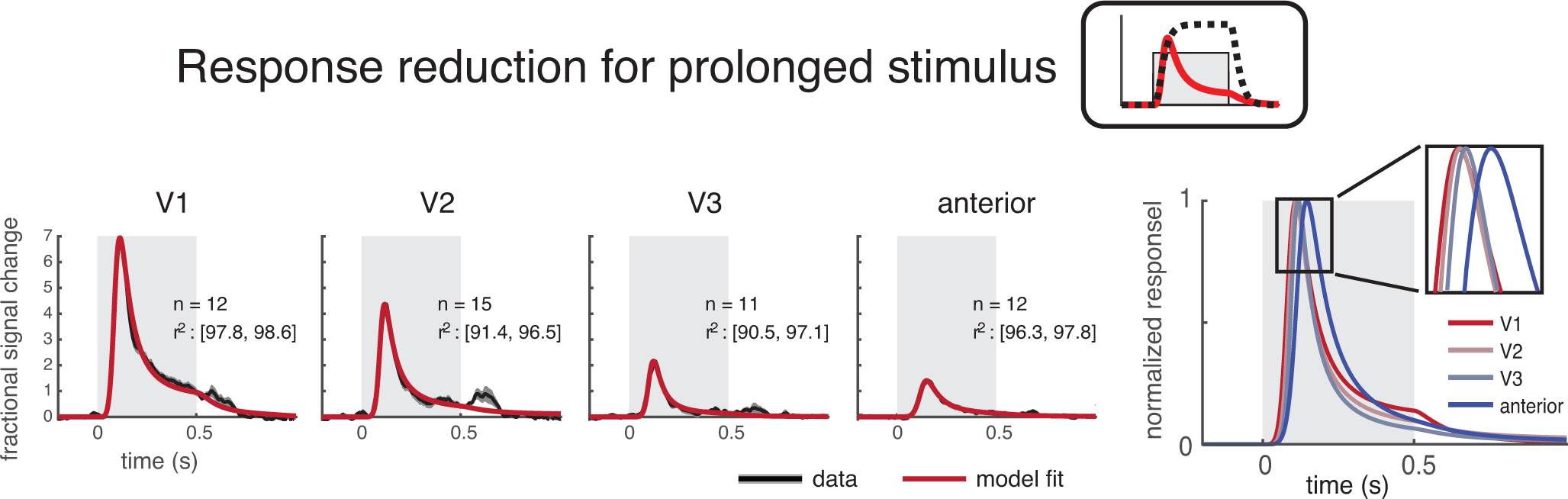
The DN model captures the response reduction for prolonged stimuli at different cortical locations. The DN model fits (red) accurately describe the ECoG broadband time course (black) in multiple ROIs. Data were averaged across trials and electrodes within ROIs, and models were fit to the average data. Each trial had a 500-ms stimulus (gray box) followed by a 500-ms blank. Plots show the mean and 50% CI for data (bootstrapped 100 times across electrodes within an ROI), and the model fit averaged across the 100 bootstraps. The number of electrodes per ROI and the 50% CI of model accuracy (r^2^ per bootstrap) are indicated in each subplot. The model fits for the 4 ROIs are plotted together on the right, scaled to unit height. For this plot, the latency was assumed to be 0 for each ROI, so that the difference in time to peak reflects a difference in integration time rather than a difference in response latency.

We derived two interpretable summary metrics to quantify model behavior in each ROI (Figure 4A): time to peak (T_peak_) and asymptotic response amplitude (R_asymp_). Each metric quantifies some aspect of the model response to a sustained stimulus. T_peak_, the model predicted response time to peak to a sustained stimulus indicates the length of the temporal summation window. T_peak_ was shortest in V1 and V2 (120-125ms), and longer in the more anterior areas (~145 ms). This summary metric excludes the onset latency, which was fit as a nuisance parameter, and hence a longer T_peak_ reflects a longer summation period, not a longer latency to respond. R_asymp_ is the ratio between the peak amplitude and the sustained amplitude. A low R_asymp_ indicates a larger extent of normalization. R_asymp_ was highest in V1 (least normalization), and decreased substantially in extrastriate areas, paralleling previous observations about non-linearities in spatial summation across visual areas (20). We summarized the differences between ROIs using these derived metrics instead of using the DN model parameters because the relationship between a single model parameter and the model output tends not to be straightforward. For example, either increasing *n* or decreasing *σ* leads to a decreased sustained response, as shown in Figure 2B, and hence neither parameter alone is a sufficient description of the amount of normalization. Although the separate model parameters are less easily interpretable, they tend to show some of the same patterns: shortest time constants in V1 and longest in the anterior maps (Figure S2).

**Figure 4.**
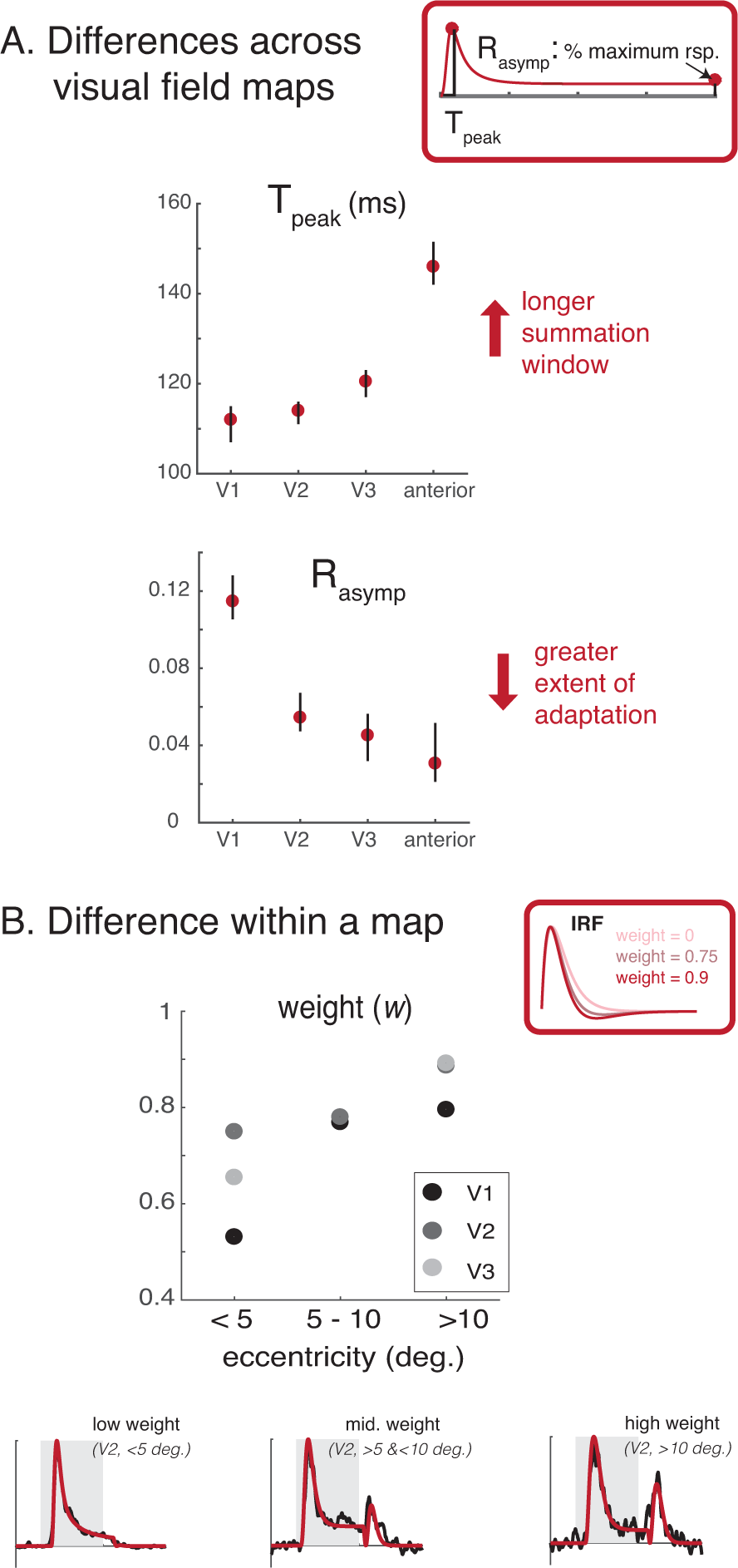
The DN model captures difference of temporal dynamics at different cortical locations. *(A) Temporal summation window length, as well as the extent of normalization increases along the visual hierarchy.* The model fits were summarized by two metrics. T_peak_ is the duration from the onset of a sustained stimulus to the peak response, excluding the onset latency. T_peak_ is longer for later ROIs, ranging from ~115 ms (V1) to ~145 ms (anterior ROIs). R_asymp_ is the level at which the response asymptotes for a sustained stimulus, as a fraction of the peak response. A smaller R_asymp_ indicates a greater extent of normalization. R_asymp_ is largest in V1 (~0.12) and declines in extrastriate areas. See Figure S3 for individual electrode results. *(B) Offset response as a function of eccentricity.* The left plots show the time series and model fits to 3 example electrodes. The offset response increases from fovea to periphery. This pattern holds across all 3 ROIs, as shown in the dot plot. Each dot is the mean weight (*w*) on the negative lobe of the biphasic response. Larger values of *w* predict larger offset responses.

### Differences as a function of eccentricity

Previous work has shown that within V1, regions with more peripheral eccentricities are more sensitive to visual transients (22). Inspection of our data in V1-V3 agrees with this pattern, as some electrodes with more peripheral receptive fields show a small positive deflection 100-200ms after stimulus offset. This offset transient was not salient in the mean time-series across electrodes (Figure 3), but it was clear in some individual electrodes (Figure 4B, Figure S3).

To quantify the offset transient response, we fit the DN model with varying *w* (weight of the negative gamma function in the IRF). For tractability of the model fit to individual electrode time series, we fixed the time to peak of the negative gamma function to be 1.5 times t (time to peak of the positive gamma function) and the exponent parameter *n* at 2. For each visual map, we separated the electrodes into three electricity bins (<5, 5-10, and >10 degrees) and averaged the parameters fitted to individual electrodes within a bin.

The model provided excellent fits to the full time-course of the response in individual electrodes including stimulus offset (Figure S3). For V1, V2, and V3, electrodes with peripheral pRF centers had higher weights (~0.8, >10 degrees) on the negative lobe of the impulse response function compared to foveal electrodes (~0.5, 0-5 degrees), consistent with fMRI studies showing that peripheral visual areas are more sensitive to stimulus transients (22, 23). We did not perform the same analysis for the more anterior areas due to an insufficient number of electrodes within each visual field map and in each eccentricity bin.

### Generalization across instruments

Above, we showed that the DN model accurately fits the ECoG broadband time series from different visual areas and different eccentricities. Here, to test generalizability, we fit the model to example time courses from 3 measurement types in early visual cortex obtained from prior publications (Figure S2A). Each time course was the response to a static contrast pattern viewed for a few hundred ms: (1) single neuron spike rates from macaque V1 (8); (2) multiunit spike rates (by taking the envelope of the bandpass filtered raw signal between 500 and 5k Hz, see method) from depth recordings in human V2/V3 (24); and (3) LFP from the same depth recordings in human V2/V3 (24). The time courses of the 3 measurements, although differing in detail, have a common pattern: there is a large, initial transient response after stimulus onset, followed by a reduction to a lower, more sustained response. This pattern was accurately fit by the DN model prediction, explaining 93% to 99% of the variance in the 3 responses. This transient/sustained pattern in these example time courses is similar to that observed in many other electrophysiological studies (e.g., 4, 5, 25).

### Phenomena 2&3: Sub-additive temporal summation and reduced responses for repeated stimuli

In our prior fMRI studies (6), we fit a static normalization model to the fMRI BOLD amplitude in response to one- and two-pulse stimuli of various durations and interstimulus intervals. Responses of the one-pulse stimuli of different durations demonstrated sub-additive temporal summation (schematic in Figure 1B), and responses to the two-pulse stimuli were consistent with reduced responses for repeated stimuli (Figure 1C). Here, we asked whether the same dynamic normalization model fit to ECoG data (previous section), with the same parameters, accurately predicts the previously published fMRI responses, thereby accounting for these two sub-additive temporal phenomena.

In brief, in the fMRI experiment subjects were presented with a large-field contrast pattern either once or twice per 4.5-s trial (Figure 5, top). For single-presentation trials, the stimulus duration varied from 0 (i.e., no stimulus) to 533 ms. For double presentations, the image was viewed twice for 134 ms each, with an inter-stimulus interval spanning 0 to 533 ms. The DN model in the previous section was fit to ECoG data with 500-ms stimuli. To use the ECoG models to predict the fMRI responses, we did the following two steps. First, for each of the 4 ROIs, we used the median model parameters across ECoG electrodes (Figure S3A) to generate broadband time-course predictions for the 13 distinct temporal stimuli used in the fMRI experiment. To convert the time-varying DN output to a predicted BOLD amplitude, we summed the DN output time-series and scaled this value to percent BOLD by applying a gain factor. Because the DN model parameters were derived from the ECoG data alone, there were no free parameters other than the gain. Although the DN models were solved with different participants, different stimuli, and a different instrument, they nonetheless accurately fit the BOLD data (*r*^2^ = 94%, Figure 5, bottom). This is more accurate than predictions from a linear model *(r*^2^ = 81%).

**Figure 5.**
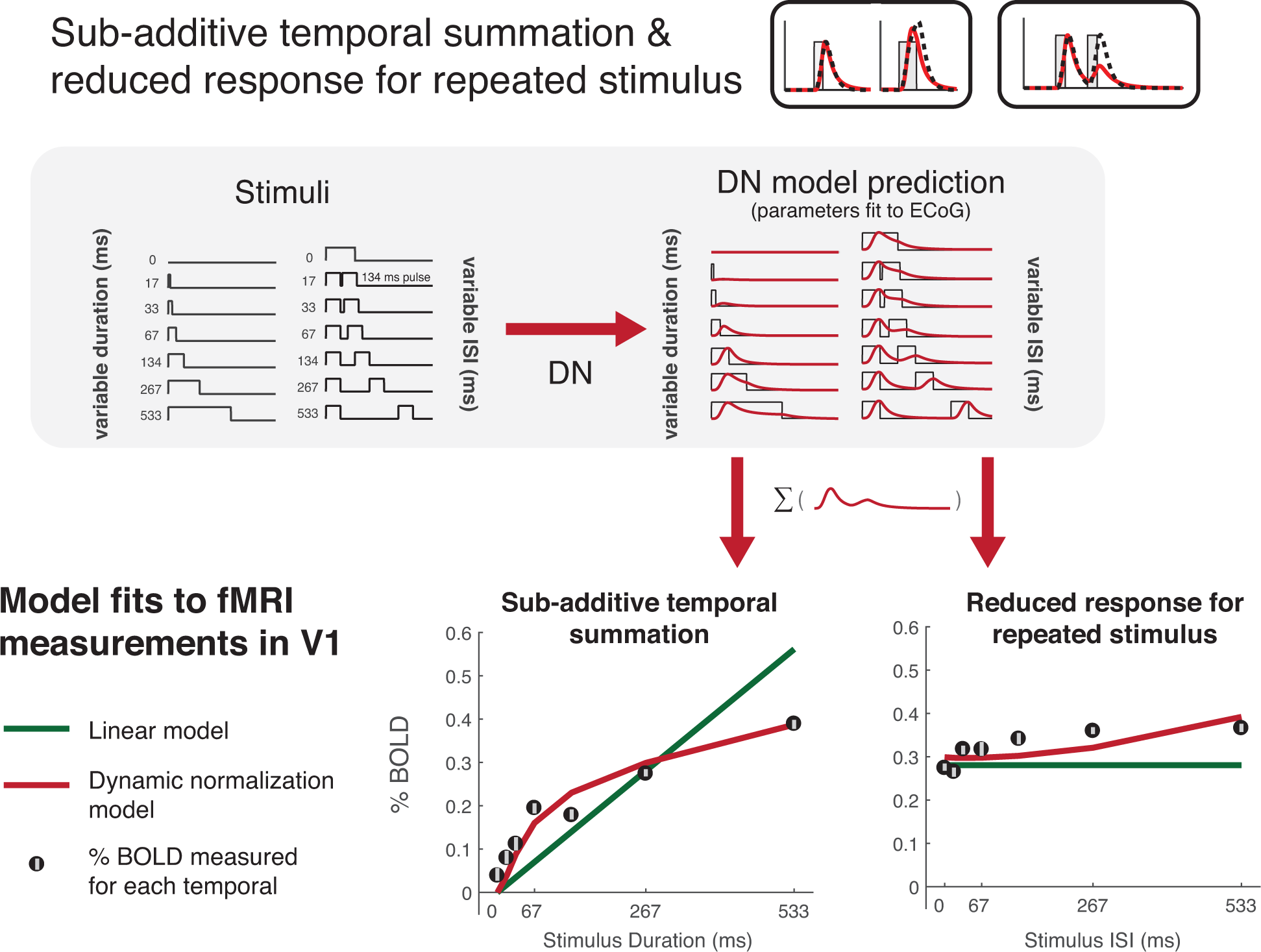
DN model captures sub-additive temporal summation and adaptation. There are two types of temporal profiles used for the fMRI experiment: one-pulse stimuli with varying durations and two-pulse stimuli (134 ms each), with varying ISI. To generate DN model predictions to these stimuli, we used the median DN parameters fit to the V1 broadband time course measured in individual electrodes (Figure S3). To convert the prediction to percent BOLD, we summed the predicted time course for each temporal profile and fit a single gain factor to minimize the difference between the predictions and the fMRI data. The DN model predictions (red) better capture the BOLD data than the linear prediction (green) (r^2^: 0.94 vs. 0.81).

Both the measured BOLD response and the predictions derived from the ECoG model fits show two patterns consistent with neuronal phenomena schematized in Figure 1. First, the BOLD signal shows evidence of sub-additive temporal summation, in that the response to long-duration stimuli are less than the linear prediction, and response to short duration stimuli are greater than the linear prediction. This pattern is accurately captured by the DN model derived from ECoG, but not from a linear model without any normalization (compare red versus green line fits in Figure 5, left plot).

Second, the bold signal shows evidence of reduced responses for repeated stimuli. This can be seen when the interstimulus interval for the two-pulse stimuli is short and the response is low (adaptation), compared to when the interval is longer and the response is higher (recovery from adaptation). This pattern is not predicted by a linear model, for which the total predicted response is the same irrespective of the interstimulus interval, but it is predicted by the DN model compare red versus green lines in Figure 5, right plot).

### Phenomenon 4: Different dynamics at low contrast, measured using single unit spike rate

In this section, we generalize the model to ask whether it can account for the response time course in single unit data from macaque visual cortex with variable stimulus contrasts. In previous sections, the input to the DN model was a time course that took only binary values, 1 whenever the stimulus was present and 0 whenever it was absent (neutral gray screen). To generalize the model to different contrast levels, we specified the model input as the stimulus spatial contrast, spanning 0 to 1, and fit this model to spike rate data from single units in macaque V1 in response to static contrast patterns (8). Other than the change in allowable inputs (continuous rather than binary), the model form itself was identical to that used to fit the ECoG broadband data.

The single unit data showed systematic differences in temporal dynamics as a function of stimulus contrast, consistent with previous reports from cat and macaque V1 (8, 15, 26). In particular, the response peak is both later and lower than the response to high contrasts (Figure 6A), similar to the schematic depicted in Figure 1D. To test whether the DN model matches this pattern, we fit the model to the response time courses for 3 complex cells, in which the stimulus contrast systematically varied between trials. For each cell, we fit a single set of model parameters for all data (10 time-courses corresponding to the 10 contrast levels). The DN model predictions behaved similarly to the data: when contrast was low, the time to peak was delayed and the peak amplitude was reduced (Figure 6B). The reduced amplitude in the model occurs because the linear response (numerator) is lower at low contrasts. The delayed time to peak occurs because there is less normalization (lower values in the denominator).

**Figure 6.**
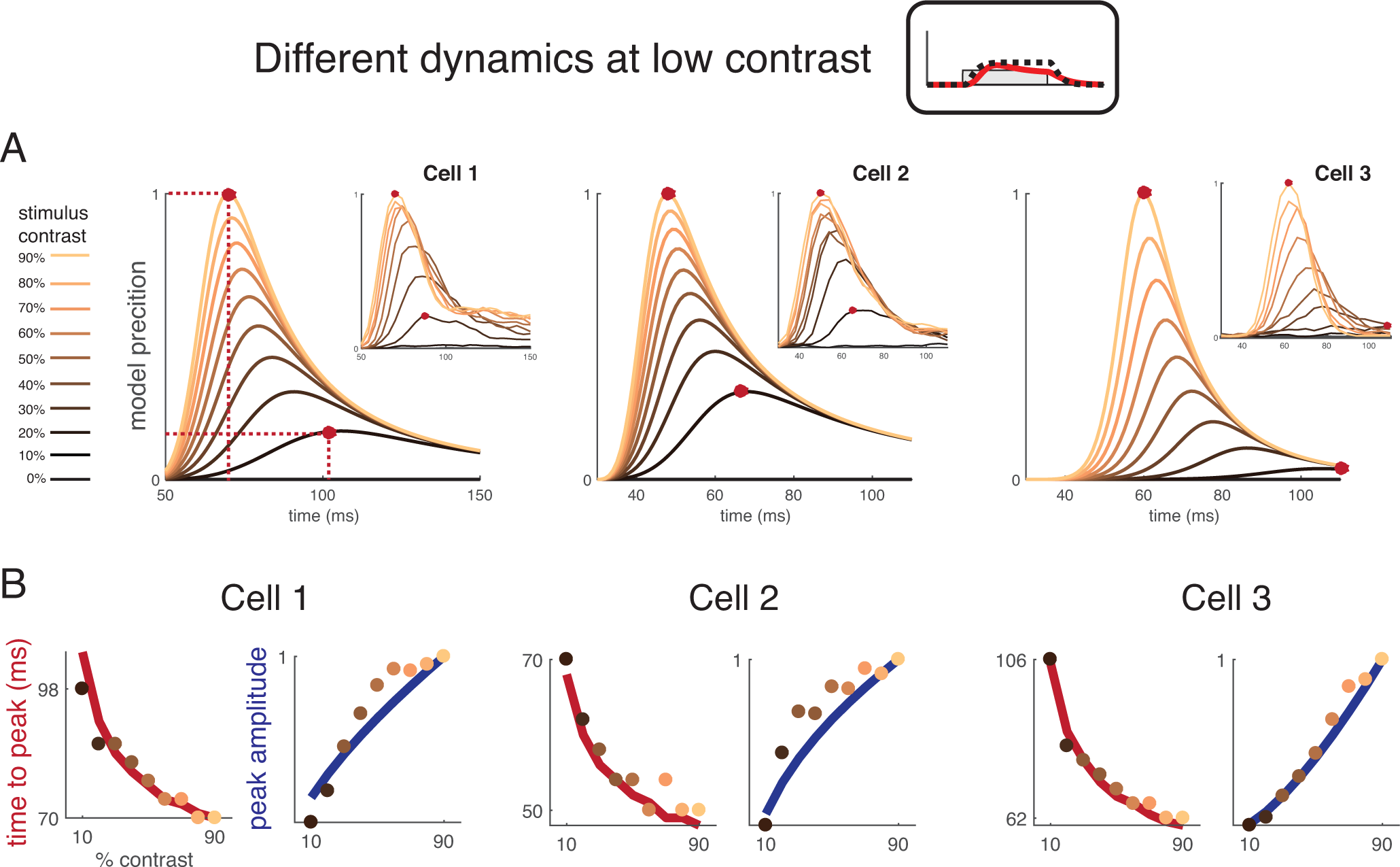
DN model captures delayed response at low contrast. *(A)* The DN model was fit to single unit spike rate data from macaque V1, with stimulus contrasts ranging from 0% to 90%. The input time course for model fitting was scaled to the stimulus contrast. A single model was fit to all stimuli (10 time-courses) separately for each of the three cells. Model fits are shown in the main plots and data in the insets. The model captures both the lower response amplitudes and slower temporal dynamics at low contrast. Data from (8), provided by W Geisler. *(B)* Time to peak (ms) and peak amplitude (normalized spike rate) for single unit data as a function of contrast. The 3 cells are those plotted in (A). The data points are the cell responses and the curved lines are the DN model fits. The colors of the dots match the colors in (A), indicating stimulus contrast.

## Discussion

We proposed a model of the temporal dynamics of neuronal responses in visual cortex. We demonstrated the generality of the model in 3 different ways. First, the model accurately accounted for diverse temporal phenomena, including sublinear summation, adaptation, and the slower dynamics at low contrast. Second, the model generalized across measurement types, including BOLD fMRI, ECoG, and single unit firing. Third, parameters of the model varied in systematic ways across cortex, both in terms of eccentricity within a map as well as between maps, enabling us to quantify regularities in temporal dynamics across cortex.

### Dynamic normalization

The model comprised 3 canonical computations: linear filtering (convolution), exponentiation (squaring, or close to squaring), and gain control (divisive normalization). A critical feature of the model that distinguishes it from our previous work (6) is that the divisive normalization is dynamic rather than static: the degree of normalization at any moment depends on the recent history of responses. As a result, the initial predicted response following stimulus onset is less normalized than the later response, causing the response to begin with a brief transient followed by a lower-level sustained response. This transient-sustained pattern is widely observed in neuronal systems and may reflect the increased importance of new information (transient component) while still conveying information about ongoing sensory inputs (sustained component). A history-dependent divisive normalization may be part of an encoding strategy to temporally decorrelate inputs, just as spatially dependent normalization can reduce correlations in the spatial image representation (27).

The dynamic normalization in the DN model differs from several other models of temporal dynamics in visual cortex, in which normalization is implemented via a change in conductance in a resistor-capacitor (RC) circuit (9, 15, 16). In these models, the conductance is determined by the instantaneous population response, and therefore the degree of normalization rises at the same rate as the linear response. In the implementation by Sit and colleagues (16), the model does not predict an onset transient, but rather a gradual rise until reaching a plateau. This accurately matches the time course of the voltage sensitive dye measurements in their study, though it differs from the time course of ECoG and spiking responses, which often show sharp onset transients (Figure 3, Figure S2A). In the two studies by Carandini and colleagues (9, 15), the stimuli were periodic and the models accurately predicted the response amplitude and phase in the steady state; the models were not tested against onsets and offsets, which might have shown the need for a dynamic (slower) form of normalization. An exception is a two stage RC circuit model of LGN responses, in which the first stage (light adaptation) depends on the recent luminance levels (11).

The presence of a delayed suppressive signal, as proposed in our DN model, does not preclude the possibility that there are also more rapid suppressive signals. In fact, both psychophysical (28) and neurophysiological studies (29) suggest that local cross-orientation suppression is rapid whereas surround suppression is sluggish. Because the stimuli used in our ECoG and fMRI experiments were large, the responses likely included effects of surround suppression. An important goal for future work will be to develop an integrated space-time model to evaluate how the spatial pattern of the stimulus affects the temporal dynamics of the responses.

### Related computational frameworks

An influential model of temporal processing in visual psychophysics proposes that the visual system encodes stimuli in a small number of temporal channels, with at least one channel that is more transient and one more sustained (17, 30, 31). A form of this model has been adapted to explain temporal dynamics measured using fMRI in V1 (22) and in extrastriate cortex (23). In these models, the transient channel contains a non-linear operator (squaring) following convolution, so that both positive and negative linear responses cause positive neural responses.

There are some similarities between the two temporal channels model and the DN model: Both include convolution with a biphasic impulse response function followed by squaring. However, the two temporal channels model does not include normalization, which is critical for explaining two of the effects we observed. First, temporal dynamics are slower at low contrast (Figure 6). This effect arises in our model as a result of normalization and is absent in a model that only includes convolution and energy (Figure S4A). Second, paired pulse experiments show that the first stimulus can affect the response to the second stimulus even with a relatively long interstimulus interval (a few hundred ms). This occurs in our model as a result of the slow normalization. In a linear / energy model an effect of the first stimulus on a second stimulus can only occur if the interstimulus interval is less than the length of the impulse response function, typically thought to be short (~100 ms).

Importantly, both our study and studies employing the two temporal channels model (22, 23) have a common finding: the periphery responds strongly at both stimulus onset and offset, whereas the fovea responds more to stimulus onset. In the two temporal channels model, this is captured by a greater weight on the transient than sustained channel; in our model, this is captured by a more biphasic impulse response function.

### Differences across cortical locations

We observed 3 systematic trends across visual cortex: (1) the temporal window length and (2) the degree of normalization increased from V1 to extrastriate areas; (3) the relative sensitivity to transients (reflected in the response to stimulus offsets) increased from fovea to periphery.

### Temporal window length

The increase in temporal window length was systematic but small, increasing by about 30% from V1 to the anterior maps just beyond V3. Qualitatively, this is similar to the increase in spatial receptive field size across the cortical hierarchy, but the differences in spatial receptive fields are larger: receptive field size more than doubles from V1 to V3, and increases by at least 4 times from V1 to V4, measured with either single units (32–35) or fMRI (20, 36, 37). Other studies have also found a hierarchy of temporal processing human and macaque cortex. Hasson et al (38) quantified temporal windows in human based on the response reliability to scrambled movie segments. They found evidence for very long temporal windows in high level areas such as the superior temporal sulcus (> 10 s). These longer windows compared to our results are likely a result of both the brain areas studied and the methods.

Temporal cortical hierarchies have also been measured (39) and modeled (40) in macaque. Murray and colleagues (39) found that the time-scale of an area while the animal was at rest (time-constant of the temporal autocorrelation function) was relatively short in early sensory areas (~100 ms or less) and longer in higher level association areas (up to ~300 ms), more commensurate with our results. In our study, we modeled all areas with the same model form, and found that the parameters changed across areas. The model could be re-expressed as a cascade, in which later areas go through more iterations than earlier areas. We show by simulation that a cascaded DN model produces a qualitatively similar pattern of results to those we observe in higher cortical areas (Figure S4B). In their model, Chaudhuri and colleagues (40) also capture the hierarchy of temporal scales, although they do not include normalization and do not account for the shape of the temporal response, such as the transient response at stimulus onset.

### Normalization

In addition to the increasing temporal summation window length, we also found an increasing extent of normalization from early to late visual areas. This gradation of adaptation levels is consistent with previous results showing that the anterior visual field maps sum more compressively in time (6). The combination of longer temporal windows and more adaptation may together cause responses in later areas to show less sensitivity (more tolerance) to changes in stimulus duration or timing, paralleling the greater tolerance for changes in stimulus size and position (20). This pattern is also consistent with the observation that activity in early visual areas tends to stay elevated for longer durations while activity in category-selective ventral temporal areas tends to decline more rapidly following stimulus onset (41).

### Stimulus offset responses

We found that temporal dynamics varied not only between maps but also within maps. Specifically, within V1-V3, peripheral response time courses measured by ECoG tended to exhibit large transient at stimulus offset. As a consequence, the peripheral responses, dominated by the onset and offset transients, are more sensitive to changes in stimulus contrast, whereas the foveal responses are more sensitive to the stimulus duration. It is likely that these differences start to emerge early in visual processing. For example, the ratio of parasol to midget cells is higher in the periphery than the fovea, contributing to higher sensitivity to transients (42). Even within a cell class, the midget ganglion cells show faster dynamics in the periphery than the fovea (43). The greater sensitivity to transients in the periphery and sustained signals in the fovea likely reflects differences in information processing across the visual field: the periphery plays an important role in exogenous attention (responding to changes in the environment), whereas the fovea is involved object recognition and appearance.

### Generalization and future directions

Our model establishes baseline performance by demonstrating explanations of several important phenomena obtained for static, large-field images over a few hundred milliseconds. This type of stimuli is well matched to many natural tasks such as scene exploration and reading, in which fixations of (mostly) static images alternate with saccades, at approximately this time scale (44). Moreover, the model serves as a valuable platform for further development to account for other stimulus manipulations and task conditions. For example, sustained attention to the stimulus (24), presence of a surround (45), non-separable spatiotemporal patterns (motion), and stimulus history of many seconds or more (46), can all affect the time course of the response.

## METHODS

### Participants

ECoG data were re-analyzed from prior work (21). As reported previously, those data were measured from 2 participants who were implanted with subdural electrodes for clinical purposes. The participants gave informed consent to participate in the study and the study was approved by the Stanford University IRB.

Functional MRI data was re-analyzed from prior work (6). As we reported previously, these data came from 6 experienced fMRI participants (2 males and 4 females, age range 21-48 years, mean age 31 years) and were collected at the Center for Brain Imaging at New York University. The experimental protocol was approved by the University Committee on Activities Involving Human Subjects at New York University, and informed written consent was obtained from all participants before the study.

### ECoG Procedure

*Preprocessing.* The data were pre-processed as in (21). In brief, electrodes that had large artifacts or epileptic activity, as identified by the neurologist, were excluded from analysis. From the remaining electrodes, we re-referenced the time series to the common average, and then down sampled the data from the recorded frequency of either 3052 or 1528 Hz to 1,000 Hz.

*Trial structure.* At the beginning of each 1-second trial, a large field (22°) noise image was randomly selected from one of 8 image classes. Several of these image classes were chosen for studying gamma oscillations in the original paper, which differs from the purpose of the current study. For this study, we analyzed data from the noise image classes only (3 of the 8 image classes): white, pink, and brown noise (amplitude spectra proportional to 1/f^0^, 1/f^1^, 1/f^2^). Noise images tend to induce a broad gamma band amplitude increase only in field potential recordings in the visual cortex, which is thought to correlate with increased spike rate and BOLD (47). Each image was presented for 500 ms followed by a 500ms blank. We analyzed data in 1200ms epochs, beginning 200 ms prior to stimulus onset and ending 500 ms after stimulus offset.

*Broadband envelope.* We computed the time varying broadband envelope in several steps, as follows. First, we band-pass filtered the time series in ten 10-Hz bins from 70 Hz to 210 Hz (70-80 Hz, 80-90 Hz; skipping 60 Hz line noise and its harmonics) using a Butterworth filter (passband ripples < 3 dB, stopband attenuation 60 dB). For time series filtered using each frequency bin, we computed its envelope as the magnitude of the analytic function (Hilbert transform). The power in field potentials declines with frequency; therefore, we normalized the envelope of each bin by subtracting the mean and dividing by the difference of the inter-quartile range, so that each envelope had a mean 0 and an inter-quartile range of 1. We then summed the 10 envelopes to derive a single, time-varying broadband envelope. (See *dn_extractBroadband.m).*

*Broadband units*. To convert the unit of the time-varying broadband to percent signal change in each electrode, we first averaged each broadband time series across epochs. We defined the first 200 ms prior to stimulus onset as the baseline period for the epoch-averaged time course, then we computed the percent signal change by dividing the entire 1200ms time course point-wise by the average of the baseline. To equalize the baseline across electrodes, we subtracted the baseline average from the entire time-course so each electrode has trial-averaged baseline 0.

*Electrode selection.* We first selected all electrodes located in identifiable visual areas based on separate retinotopy scans. Among these location-identifiable electrodes, we only chose the electrodes that satisfy the following two criteria for further analysis: 1. electrodes whose trial-averaged broadband response during the stimulus on period (500 ms) is greater than the baseline period on average; 2. electrodes whose maximal trial-averaged broadband response is greater than 150% of the prenormalized baseline average (see *Broadband units).* (See *dn_chooseElectrodes.m)*

*Foveal versus peripheral electrodes.* Based on the retinotopy analysis, we separated the electrodes within V1-V3 into three eccentricity bins (<5, 5-10, >10 degrees) based on their estimated receptive field centers.

### Single- and Multi-unit Procedure

*Single-units preprocessing.* We re-analyzed two macaque single-unit data sets from Albrecht et al. 2002. The first data set consists of trial-averaged PSTH from 12 complex cells in V1 (Figure S2A), and each PSTH represents a distinct type of response shape (Albrecht et al. 2002, their figure 4). The stimulus for this data set is a large field spatial grating presented over 10 different contrasts and 9 spatial phases. Each PSTH in the data set is the response averaged across 40 repetitions of each pair of contrast and spatial phase combination. To generate the average single-unit time course in Figure S2 (panel A), we first duplicated each PSTH *n* times, with *n* being the number of cells within each shape category (see Albrecht et al. 2002 figure 4). We then bootstrapped over this expanded cell set by randomly sampling (100 times) with replacement. Finally, the DN model was fitted to the average of the bootstraps (see DN Model Fit).

The second data set (Albrecht et al. 2002 figure 1) consists of three cells’ responses to a 200 ms presentation of a large field stationary grating (8 different spatial phases) at 10 linearly spaced (0-90%) contrast levels (Figure 6).

*Multi-units preprocessing.* We re-analyzed the data correspond to the “Contextual modulation” experiment in (24). The stimulus used in the experiment consists of one (stationary) spatial grating restricted to a circular patch, and the rest of the screen is filled with another grating that is of the same spatial frequency, but the same or different orientation and phase. Each grating is presented at 80% Michelson contrast, and with a spatial frequency 1 cycle/degree. In each trial, the stimulus is on for 500ms before the screen returned to neutral gray. For our purpose, we pooled signals across all trial types (Figure S2A).

MUA and LFP extractions are exactly the same as described in (24). We fitted the DN model to the trial-averaged MUA signals, and the time-varying broadband envelopes within the LFP signal. The broadband extraction process of the LFP signal is the same as that described in *Broadband envelope* under the ECoG Procedure, except for one minor difference: 1. We band-pass filtered the time series in ten 10-Hz bins from 85 Hz to 175 Hz (excluding harmonics of 50Hz line noise), instead of from 70 to 210 Hz as in the ECoG data. We chose a higher starting frequency (85 Hz) here because grating stimuli tend to induce an oscillatory signal within 30-80 Hz range measured on the visual cortex, and we chose a lower ending frequency (175 Hz) because the sampling rate here is lower than the ECoG data. (See *dn_analyzeMultiData Types. m)*

### The temporal pRF Models

#### Models

*Dynamic Normalization (DN) model.* Here we introduced a temporal encoding model - the Dynamic Normalization (DN) model, to capture neuronal responses over time measured using different methods. The model takes the time course of a spatially uniform contrast pattern as input *(T*input), and produces a predicted neuronal response time course as output.

The DN model captures neuronal dynamics over time with a divisive normalization. The numerator of the model consists of a linear response: an impulse response function (IRF) *hi* convolves with a stimulus time course *S(t)* (with 1 represents stimulus on and 0 for stimulus off).

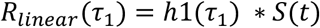

The impulse response function is represented by the weighted difference between two gamma functions:

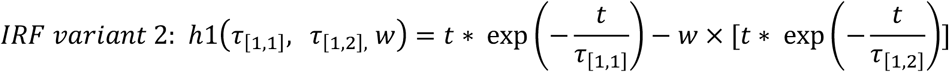

We vary *w* only in figure 4B, S3, and S4A because the ECoG time courses averaged across electrodes, as well as the spiking time courses do not show a prominent offset transient response. When *w* is varied, we fixed *n* = 2, so that the output responses would consists only of real numbers. For Figure. 3 and Figure 4A, we fixed *w* to be 0, so that a single gamma function is used as the impulse response function.

The denominator of the model is the sum of two terms, a semi-saturation constant (σ) and an exponentially filtered (low-pass) linear response. The rate of the exponential decay is determined by a parameter t2.

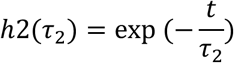

All three terms (one in the numerator, two in the denominator), are raised to the power *n*, assumed to be greater than 1.

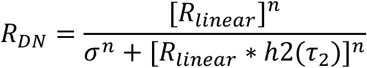

To fit the first DN model to the time series data (SUA, MUA, LFP broadband and ECoG broadband), we vary all four model parameters (*τ*_1_, *τ*_2_, *n, σ)* together with two nuisance parameters. The first nuisance parameter represents a delay that accounts for the time lapsed between stimulus onset and response onset; the second nuisance parameter scales the predicted model output to the same range as the measured signals.

To fit the second DN model to the time varying broadband measured using ECoG, for tractability we fixed the time constant of the second gamma function in the IRF to be 1.5 times of the first, i.e. *τ*[_1×2_] *= 1.5τ*_[1,1]_. We varied the weight parameter *w*, which represents the relative weight of the second (negative) gamma function in the IRF. Higher *w* links to a higher predicted post-stimulus transient response for a sustained stimulus. We further fixed the normalization parameter *n* to be 2, so that the predicted response consists of real numbers. We fit the rest of the model parameters *(τ_1_, τ_2_*, *σ)* and the nuisance parameters as in other case. (See *dn_DNmodel.m.)*

*Two temporal channels model* (Figure S4, Horiguchi et al. 2009). The model consists of a weighted sum of two components - each component is interpreted as the output of a sustained or a transient temporal channel. The output of the sustained component follows a linear computation, and the output of the transient channel follows a sub-linear computation:

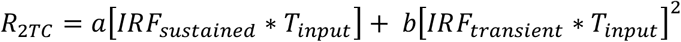

Higher weight of the transient component leads to a higher degree of the offset transient response, and higher weight of the sustained component leads to a higher level of the sustained response. (See *dn_2Chansmodel. m)*

*Compressive Temporal Summation* (Figure S4, (6)). The compressive temporal summation (CTS) model is similar to the DN model except that the normalization is instantaneous instead of delayed as in the DN.

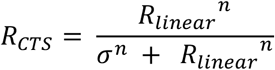

For simplicity, we assumed n = 2 when fitting this model to the ECoG time course in Figure S4. (See *dn_simpNormModel. m)*

*Cascaded DN model* (Figure S4). When fitting the DN model to the ECoG broadband time courses, we assumed that for each visual area, the DN model takes a stimulus as input and produces a response time course as output. Alternatively, we also illustrate the behavior of a two-stage cascade model: we used the output of the V1 model as the input to a second, identical model, and show that this produces responses qualitatively similar to V3AB.

#### Parameter estimation

*DN model for ECoG.* We used a two-stage approach to fitting the DN model, first to obtain seeds (grid fit) and then to estimate parameters (search fit). For the grid fit, we computed model predictions to the 500 ms stimulus for 10,000 combinations of *τ*1, *τ* 2, n, and σ (*τ*_1_: [0.07, 1], *τ*_2_: [0.07, 1], n: [1, 6], σ: [0.01, 0.5], each parameter in the range with 10 equal steps). Using linear regression on the data time course, we derived the gain factor, g, and the variance explained for each of the 1,000 predicted time series. For each bootstrapped response time course in each ROI, the set of parameters that generated the highest variance explained was used as the seed for the search fit. (See *dn_gridFit.m)*

For the search fit, we used a bounded nonlinear search algorithm in Matlab *(fminsearchbnd.m)*, run once per ROI per bootstrap. The search finds the parameters that minimize the squared error between the predicted and the measured time broadband course. The lower bound used for the search fit was [0.07, 0.07, 1, 0.01, 0.0001] for *τ*_1_, *τ*_2_, n, σ and a shift parameter that accounts for the delay between stimulus onset and response onset. The upper bound used for the search fit was [1, 1, 6, 0.5, 0.1]. In principle, the delay parameter is important, since the time at which the signal from the stimulus reaches cortex is delayed, and the delay varies across visual field maps, and could be as high as 50-150 ms. However, the impulse response function includes a slow ramp, and the broadband envelope extraction contains a small amount of blur. Hence in practice, the shifts were quite small (< 10 ms), and not informative about the latency of neuronal response. To summarize the fit, we plotted the mean of the predicted time course across bootstraps and the standard deviation at each time point as the confidence interval. (See *dn_fineFit.m)*

*DN model for single-unit data with variable stimulus contrast (Figure 6).* We fit the DN model to three cells’ response time course to a 200-ms stimulus contrast increment at 10 different contrast levels (0% - 90% contrast with 10 steps of equal increment). We fit one set of DN parameters to all 10 response time courses for each cell by minimizing the squared error between the data and prediction. We seeded the search fit for the first two cells with [0.1, 0.1, 2, 0.2, 0.03], and the last cell with [0.1, 0.1, 3, 0.1, 0.04] (*τ*_1_, *τ*_2_, n, σ, and a shift parameter). Then we used *fminsearch.m* in Matlab for the search fit. (See *dn_mkFigure_fitDN2ContrastSUA.m)*

*DN model for fMRI BOLD amplitude (Figure 5).* To predict the fMRI response from the DN model, we used the parameters fitted from ECoG data for each electrode (Figure S3), took the median of each parameter within each ROI, and generated a neuronal time course for each of the 13 distinct temporal profiles from the fMRI experiment. Then we summed each predicted time course, and finally scaled the sum by a gain factor, g. The only free parameter was the gain factor. (See *dn_fitDNECoG2fMRI.m)*

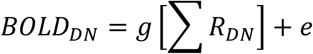

*Biphasic DN model fit to ECoG broadband (Figure 4B).* To fit the DN model with a biphasic IRF to the broadband time course estimated for individual ECoG electrodes, we varied five model parameters together: *τ*_1_ *w, τ*_2_, σ, and a shift parameter. “*w*” is the weight of the negative pulse in the IRF. The length of the second pulse, *τ*_2_, was assumed to be 1.5 times *τ*_1_ and *n* was assumed to be 2. For each electrode, we generated four predictions from these four sets of parameters first: [0.02, 0.8, 0.15, 0.1, 0.05]; [0.03, 0.8, 0.1, 0.2, 0.05]; [0.02, 0.4, 0.15, 0.1, 0.05]; [0.03, 0.4, 0.1, 0.2, 0.05]. These parameter sets differ in the extent of normalization and the extent of the post-stimulus transient response. We picked the parameter set that generated the highest variance explained for each electrode, and used the set as the seed for a further search fit. (See *dn_mkFigure_bidnFit2ECoG.m)*

#### Model accuracy

Throughout the paper, we summarized model accuracy as the variance explained, *r*^2^, the square of the Pearson-correlation coefficient *r*.

### Public Data Sets and Software Code

To ensure that our computational methods are reproducible, all data and all software will be made publicly available via an open science framework site, https://osf.io/z7e3t/. The software repository will include scripts of the form *dn_mkFigure2.m* to reproduce figure 2, etc., as in prior publications (6).

## Acknowledgements

We thank Dora Hermes for helpful discussion and for helping us analyze ECoG data from prior work. We also thank Josef Parvizi and the Stanford Human Intracranial Cognitive Electrophysiology Program for helping us with ECoG data acquisition for a prior paper, which was re-analyzed for this paper. We thank Bill Geisler for providing single unit data. We thank David Heeger, Mike Landy, and Jennifer M. Yoon for comments on an earlier draft of this manuscript. The research was supported by NIH grants R00-EY022116 and R01-MH111417.

## Supplementary Material

**Figure S1.**
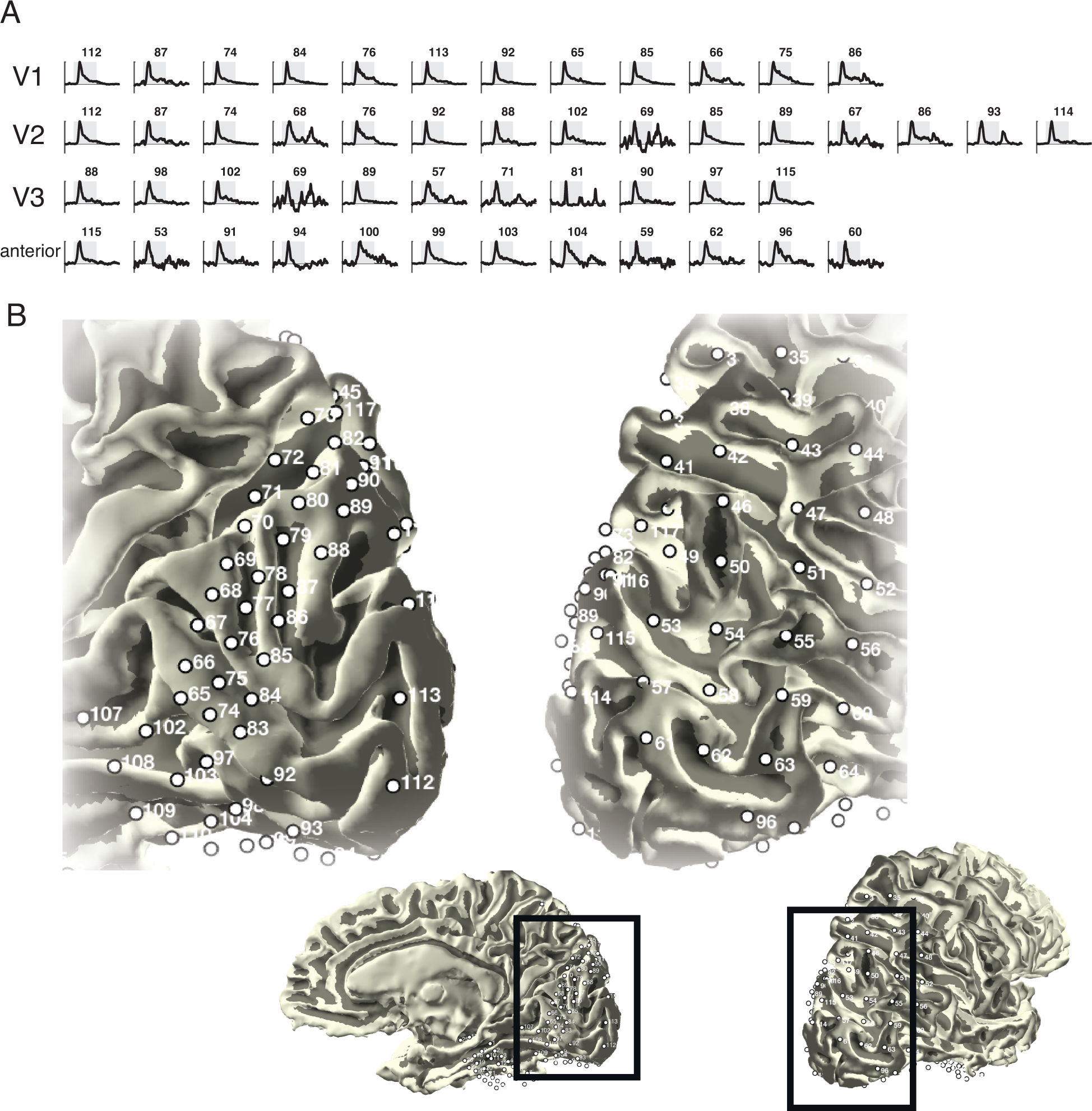
Individual electrode responses. The plots show the ECoG broadband time course in individual electrodes from ECoG subject S1, averaged across 90 trials (30 repeats each of three stimulus types). Each row shows electrodes from one ROI. Some electrodes (e.g., 74) are in two rows, since the electrode was near an ROI boundary. The plots are color coded by eccentricity bin (0-5°, 5-10, >10°). The pRF location was based on a separate ECoG pRF data set published previously (Winawer et al., 2013). The two mesh images show a magnified view of Si’s right occipital lobe, exposing the medial surface (left) and lateral surface (right). Insets show the zoomed-out view of the cortical mesh.

**Figure S2.**
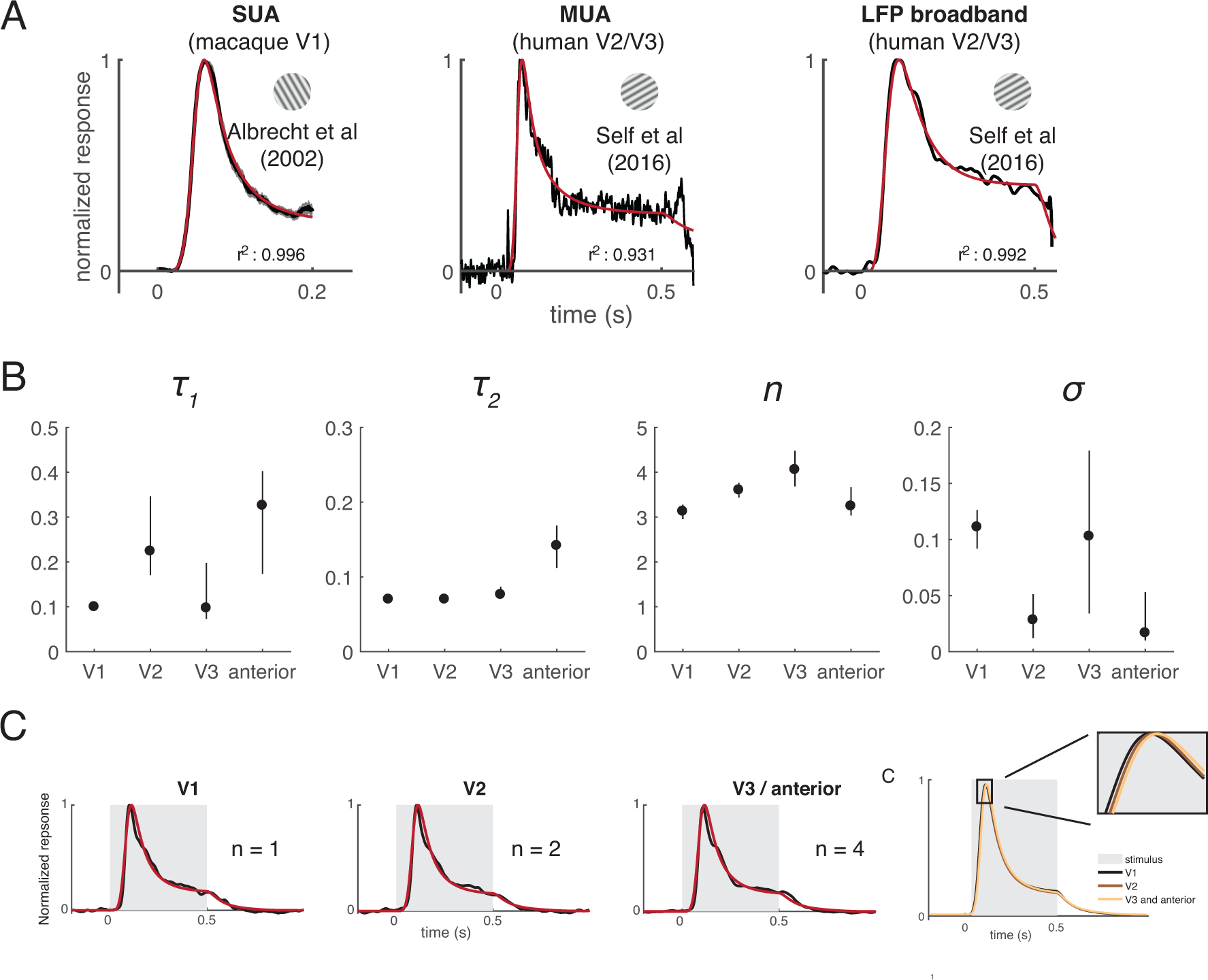
Response reduction for prolonged stimuli. (A) Response time courses from 3 different recording methods are shown. In each plot, the data are in black (±1 sem in gray) and the DN model fit in red. Left: single unit spike rates, averaged across neurons in macaque V1. Middle: Multiunit spike rates from human V2/V3. Right: High frequency broadband power (LFP) from human V2/V3. (B) DN model parameters from human ECoG. The model parameters in each of 4 ROIs are shown for the data plotted in the main text (Figure 3A). (C) ECoG broadband responses in 3 ROIs from subject S2. Plotting conventions as in Figure 3.

**Figure S3.**
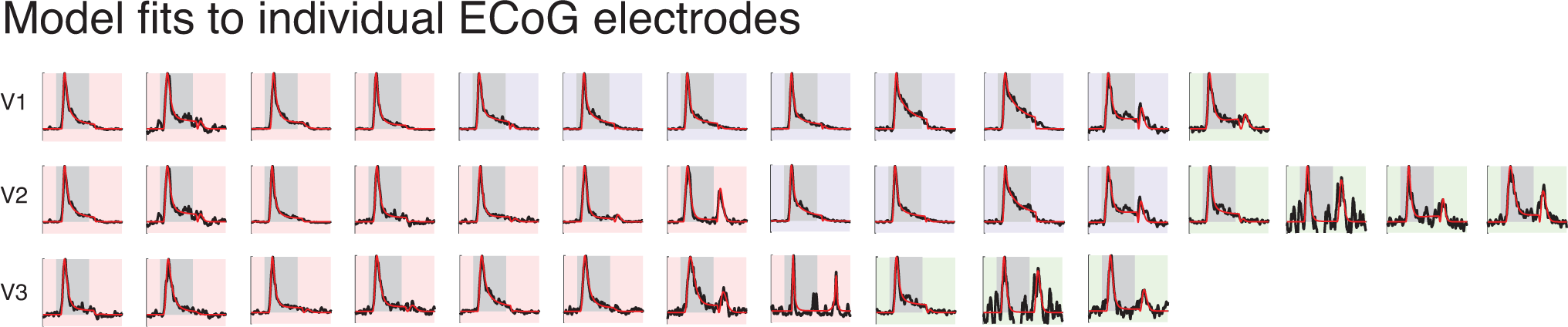
Effects of eccentricity and contrast on temporal dynamics. Individual electrode time courses and DN model fits in V1-V3. The background color indicates the eccentricity bins: 0°-5° (red), 5°-10° (purple), and >10° (green). There is a general tendency toward greater offset responses in more peripheral electrodes.

**Figure S4.**
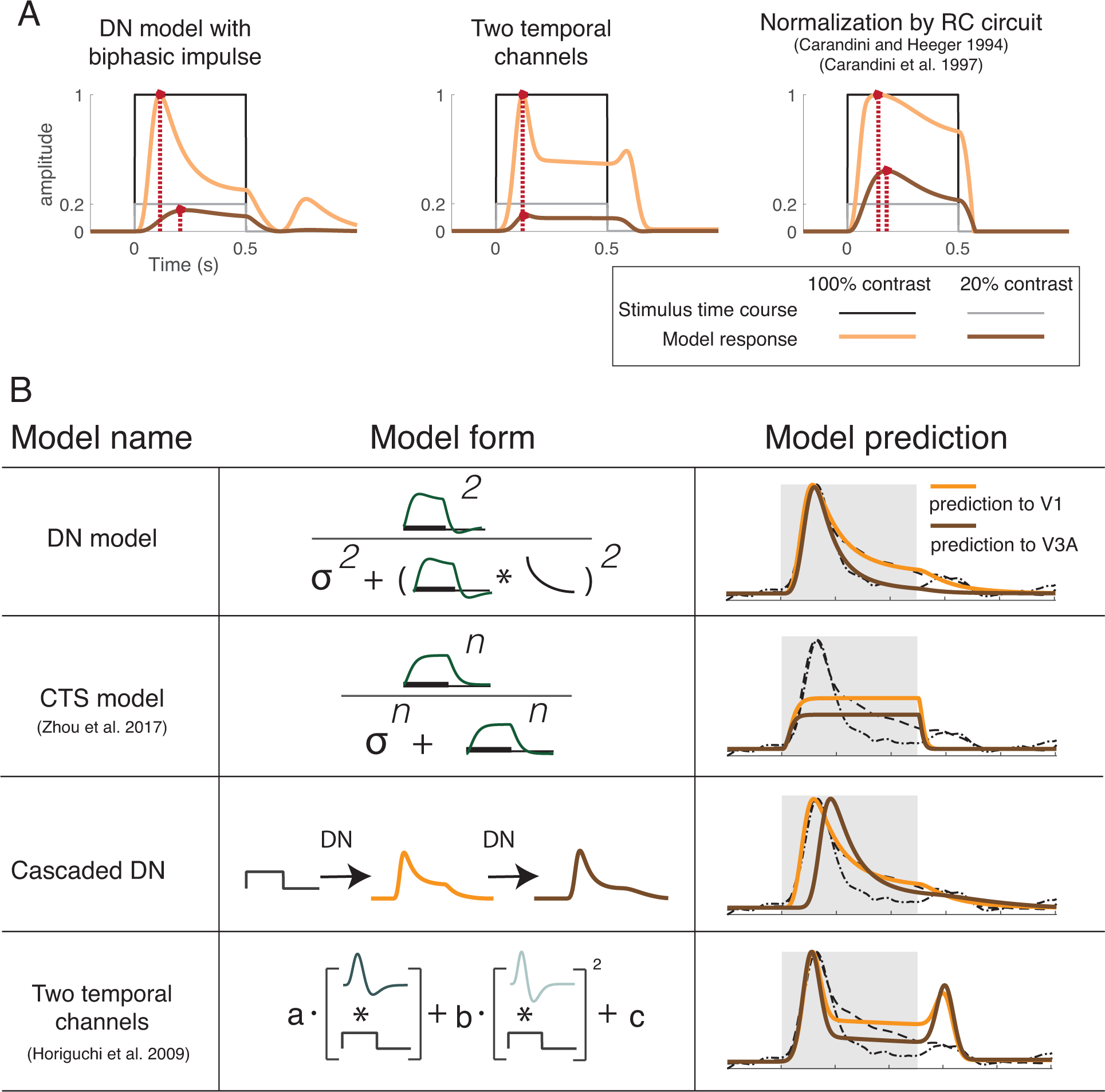
Model comparisons. (A) Each plot shows the predicted responses from one model for 500-ms stimuli at full contrast (orange curve) or 20% contrast (brown curve). The dark lines indicate the stimulus time course. The red dots show the peak response. The DN and RC circuit models, but not the two temporal channels model, show slower dynamics at low contrast. The DN and two temporal channels model but not the RC circuit model show sharp transients at stimulus onset. (B) Each row shows fits to ECoG data from one model. Data are fit to ECoG broadband time courses from one electrode in V1 and one electrode in V3A. The two ECoG time courses are shown by the gray dashed lines in the third column. These are the same in every row. The model fits are shown in orange (V1) and brown (V3A). The difference in the first two rows show the importance of the low-pass filter for normalization (DN model but not CTS model). Without it, the predicted response does not show a transient / sustained pattern. The third row shows an alternate method for achieving longer integration and more normalization in later areas: a cascade of identical DN models rather than a single model with different parameters. See Methods for model details.

